# Sequence-based Optimized Chaos Game Representation and Deep Learning for Peptide/Protein Classification

**DOI:** 10.1101/2022.09.10.507145

**Authors:** Beibei Huang, Eric Zhang, Rajan Chaudhari, Heiko Gimperlein

**Affiliations:** Department of Experimental Therapeutics, The University of Texas MD Anderson Cancer Center, Houston, TX 77054, USA; Eurofins Beacon Discovery, San Diego CA 92121, USA; Maxwell Institute for Mathematical Sciences and Department of Mathematics, Heriot–Watt University, Edinburgh, EH14 4AS, UK

## Abstract

As an effective graphical representation method for 1D sequence (e.g., text), Chaos Game Representation (CGR) has been frequently combined with deep learning (DL) for biological analysis. In this study, we developed a unique approach to encode peptide/protein sequences into CGR images for classification. To this end, we designed a novel energy function and enhanced the encoder quality by constructing a Supervised Autoencoders (SAE) neural network. CGR was used to represent the amino acid sequences and such representation was optimized based on the latent variables with SAE. To assess the effectiveness of our new representation scheme, we further employed convolutional neural network (CNN) to build models to study hemolytic/non-hemolytic peptides and the susceptibility/resistance of HIV protease mutants to approved drugs. Comparisons were also conducted with other published methods, and our approach demonstrated superior performance.

**Supplementary information:** available online

## 1 Introduction

Chaos Game Representation (CGR) is an effective graphical representation method for 1D sequences including text, genetic codes, and protein sequences, thus it has attracted significant attention over the years (Olyaee, Khanteymoori, and Khalifeh 2019; Basu et al. 1997; Joseph and Sasikumar 2006). As a powerful tool, CGR is usually used to describe the attributes of biological sequences and predict their certain functions through deep learning (DL) (Heider, Verheyen, and Hoffmann 2011; Ge et al. 2019; Lochel et al. 2020), which has also been successfully used in the field of protein structure modeling (Jumper et al. 2021; Baek et al. 2021). Additionally, a variety of techniques, such as variational autoencoder (VAE) (Hawkins-Hooker et al. 2021; Ding, Zou, and Brooks 2019) and supervised autoencoders (SAE) (Le, Patterson, and White 2018), have been developed to extract crucial features from amino acid sequences. For instance, the distribution of VAE encoded data points in their latent space is close to a standard normal distribution and these models are capable of extracting the latent subspace features for interpretation (Chen et al. 2016; Klys, Snell, and Zemel 2018). On the other hand, SAEs provide us with options to building deep generative models which can project the original data points into latent representations that benefits the desired tasks such as classification. In other words, SAE is an autoencoder with addition of a supervised loss at the representation layer, and such an extra term should better guide the learning toward data representations that are conducive to the task (e.g., classification). Most recently He et al. developed CODE-AE based on a SAE scheme and constructed a feature encoding module that can be easily tuned to adapt to the different down-stream tasks (He et al. 2022).

In our present study, we adopted the SAE strategy to encode peptide/protein sequences by combining with optimized CGR representations for functional classification tasks. In CGR, one of the most important elements is scaling factor (SF), which determines how the representation is generated. Hence, it is crucial to optimize SF in CGR (Olyaee, Khanteymoori, and Khalifeh 2019; Lochel et al. 2020). Herein, we attempted to achieve this by devising a new energy function to improve the encoder quality, with two approaches: **1.** Directly perform gradient descent on SF to obtain the optimized CGR images; **2.** Construct a SAE to optimize CGR representations. Furthermore, we analyzed the relation between the numerical CGR representation and the SAE encoded representation and found that they are equivalent in the latent space. Based on this observation, we employed the optimized SAE encoder to convert amino acid sequences into a new form of images that are more conducive to classification problems.

To further evaluate our approaches, we conducted two binary classifications studies: one is to determine whether a peptide is hemolytic, and the other is to model if an HIV mutant virus is susceptible to the Food and Drug Administration (FDA) approved HIV drugs. For the first study, we focused on therapeutic peptides which play a notable role in biomedical research and disease treatment (McGregor 2008; Lau and Dunn 2018; Muttenthaler et al. 2021; Drucker 2020; Fox 2013). It is known that many peptides are hemolytic and cause destruction of red blood cells, thus leading to hemolytic anemia (Fosgerau and Hoffmann 2015). The optimization of peptide hemolytic properties is an essential but time-consuming and labor-intensive task, therefore, it is crucial if we can efficiently predict the hemolytic nature of peptides during lead optimization (Chaudhary et al. 2016). It is worth mentioning that a variety of reports have been published over the years for peptide hemolytic property predictions (Raghava et al. 1994; Singh, Ansari, and Raghava 2013; Chaudhary et al. 2016). Herein, we concentrated on the optimization of the amino acid sequence representation, i.e., how to encode sequences into images to best predict their hemolytic effects in the deep learning framework. For the second classification task, we modeled the susceptibility of HIV protease (HIVP) mutants to five approved protease inhibitors: indinavir (IDV), saquinavir (SQV), nelfinavir (NFV), amprenavir (APV), and lopinavir (LPV) (Lochel et al. 2020; Rhee et al. 2006). Because of its high mutation rate, HIV virus can rapidly evolve and lead to drug resistance (Condra et al. 1995; Loeb et al. 1989), and most recently researchers uncovered a highly transmissible and damaging variant of HIV that has been circulating in the Netherlands for decades (Wymant et al. 2022). Therefore, to provide the best treatment options, it is critical if we can predict whether a patient with a specific strain of virus is sensitive/resistant to what drugs based on the individual HIVP mutation profile.

## 2 CHAOS GAME REPRESENTATION AND SUPERVISED AUTOENCODER

### 2.1. Chaos Game Representation (CGR) for Amino Acid Sequences

Although 3D structure-based methods have significantly improved peptide/protein design [21], sequence-based encodings are more versatile since they are based only on the amino acid composition and many newly designed peptides or proteins have no 3D structure available. Herein, we adopted a sequence-based encoding method to represent amino acid sequences. As one of the most typical sequence-based encoding strategies, the n-gram method counts the occurrences of n-mers (n consecutive letters, a.k.a., motifs) for a size m alphabets, leading to a *m^n^* dimensional representation of the initial sequences. For peptides and proteins, *m* = 20 corresponds to the number of common amino acids. There are two ways to define n:

1. **as a constant,** for example, n=3. In this way, all 3-mers (motifs) form a set that contains 20 × 20 × 20 elements. Practically, we count the occurrences of the motifs in the sequences, and the whole sequence is represented by a list of motifs.
2. **as a variable,** determined based on classification of the observed sequences, e.g., as defined by **MERCI** (Motif EmeRging and with Classes Identification) (Vens, Rosso, and Danchin 2011). Since some conserved motifs have different lengths (different n) in sequences, this method not only more accurately represents the biological functions of sequences, but also more efficiently characterizes them for classification.

To demonstrate the concept of CGR, **Fig. 1** shows the transformation process by using randomly selected 50 peptides from our curated hemolytic datasets (described in **Section 3**). Among them 25 are experimentally validated as hemolytic peptides (positive annotated with 1) and the other 25 are non-hemolytic (negative annotated with 0). The analysis of these sequences obtained a set of 130 motifs (each occurred at least once), which are defined as our motif dictionary. These motifs are projected and evenly distributed in 1D axis with coordinates *P_k_*, *k* = 0,1,2 …129, as shown in **Fig. 1a**, and they are defined as the anchor points. Then each sequence, depending on the motifs contained, is represented by a series of points on x-axis as illustrated in **Fig. 1b**, with coordinates of x_1_ x_2_,… x_n_, each corresponding to the position of the respective motif. These coordinates can be derived through a recurrent iterative function (RIF) as described in **Eq. 1**:

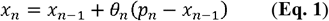

where *p_n_* denotes the position (the coordinate) of the anchor point in the dictionary corresponding to the n-th motif in the sequence; *θ* is the coefficient scaling the distance between the actual motif position *x*_*n*-1_ and the target anchor point *p_n_*, thus also known as the aforementioned scaling factor. *x*_0_ is the origin of the x-axis. Most studies with CGR for protein sequences have been using a constant *θ* = 0.5; however, this can lead to noisy images when *n* > 4 (Lochel et al. 2020). Therefore, we used different SFs to encode sequences and perform SF optimization to build the best classification models. Taking “GIMSLFKGVLKTAGKHVAGSLVDQL KCKITGGC” and “FEQTGGPDLTTGSGKRTKSDRVE HKHASQ” as examples, we first evenly distributed the 130 motifs as the anchor points over the interval [−5,5] (**Fig. 1a**). In both peptide sequences, we found 5 motifs, which are projected to the x-axis with a set of SFs using **Eq. 1** to derive the corresponding 5 encoded points (red in **Fig. 1b**).

**Fig. 1.**
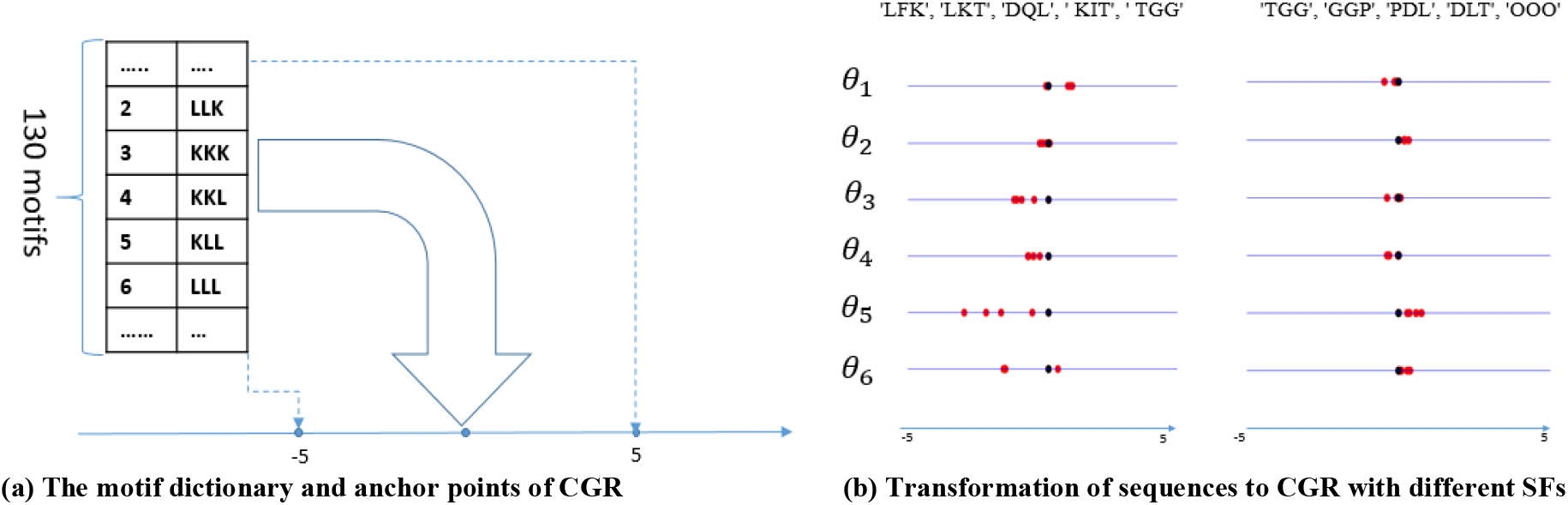
**(a).** An example of the dictionary with 130 motifs and their corresponding anchor points based on 50 peptides randomly selected from HemoPI-1. The anchor points are evenly distributed on 1D axis, denoted as *P_k_, k* = 0,1,2 … 129, with values between −5 and +5. **(b).** The first peptide sequence “GIMSLFKGVLKTAGKHVAGSLVDQLKCKITGGC” has 5 motifs, ‘LFK’, ‘LKT, ‘DQL’, ‘ KIT’, ‘TGG’, and the red points correspond to these 5 motifs encoded via **Eq. 1**, using 6 sets of SFs θ_1_, i=1…6. The coordinates of these points representing the sequence are derived based on the corresponding SF and are plotted along the respective lines. The black point denotes the respective origins xø. Using the same process, the coordinates are obtained to represent another peptide sequence “FEQTGGPDLTTGSGKRTKSDRVEHKHASQ” (on the right), for the corresponding 5 motifs ‘TGG’, ‘GGP’, ‘PDL’, ‘DLT’, ‘OOO’, based on the same 6 sets of SFs.

Of note, the “OOO” motif is a pseudo motif intentionally added when less than 5 motifs from the dictionary are found in a sequence. On the other hand, if more than 5 motifs are found, we will consider the 5 motifs that have the highest ranks in our motif dictionary. The initial values of SF were randomly generated and **Fig. 1b** shows 6 sets of SFs used during the optimization to encode the two sequences. In this way, we converted each peptide sequence to a group of points distributed on the x-axis. Presumably, if the sequences belong to the same functional class (e.g., hemolytic peptides), their corresponding encoded points should have similar spatial distributions and they should be clustered together (short distance from each other) based on the CGR representation. This lays the foundation for our optimization process. To this end, we define G(α) as the distribution of the encoded points of the sequence α on x-axis. Similarly we define G(β) for the sequence β. The difference between sequences α and β is defined as their distribution distance calculated with Jensen-Shannon Divergence *D*(*G*(α), *G*(β)). Herein, we employed point-to-origin distance as the distribution function, as described in **Appendix 1** in Supplementary Materials. Furthermore, we found that if two sequences are projected into 2D space, i.e., represented by greyscale or color images, their distance can be similarly calculated with Jensen-Shannon Divergence, as described in **Appendix 2** in **Supplementary Materials**. Based on these findings, we designed an energy function *L_CGR_*(*θ_k_*), which represents the sum of pair-wise distances of all sequences in terms of the scaling factor *θ_k_*

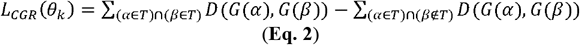

where *T* denotes one class, and (*α* ∈ *T*) ∩ (*β* ∈ *T*) means that the sequences *α* and *β* belong to the same class, while (*α* ∈ *T*) ∩ (*β* ∉; *T*) or (*α* ∉ *T*) ∩ (*β* ∈ *T*) indicates *α* and *β* are of different classes.

For **Eq. 2**, there are 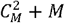 pairs of sequences to be calculated for the energy of the system (*L_CGR_*) where *M* denotes the total number of sequences. Based on our assumption, the distribution distance *D* of the same class of sequences should be smaller than that of different classes.

Such properties can be used to minimize the energy by optimizing the scaling factor *θ_k_* via

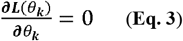

Practically, the optimization process on the whole dataset can be approximated by the batch gradient descent method. The optimized 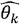 can be utilized to encode the sequences for the best representation. The algorithm to optimize SFs is illustrated in **Fig. 2(a)**, namely **Algorithm A**. The original amino acid sequences with the initial random SFs are fed into RIF (**Eq. 1**) in batch, and then the total energy *L_CGR_* and its gradients are computed accordingly. The procedures are repeated until SFs converge. We observed that the calculation of the gradient of *L_CGR_* is time-consuming and it is difficult to control the step rate (or learning rate), especially when the the number of sequences is large. Therefore, this method is not preferred in such situation. Instead, we have developed a deep learning-based method to address the issue, as described next.

**Fig. 2.**
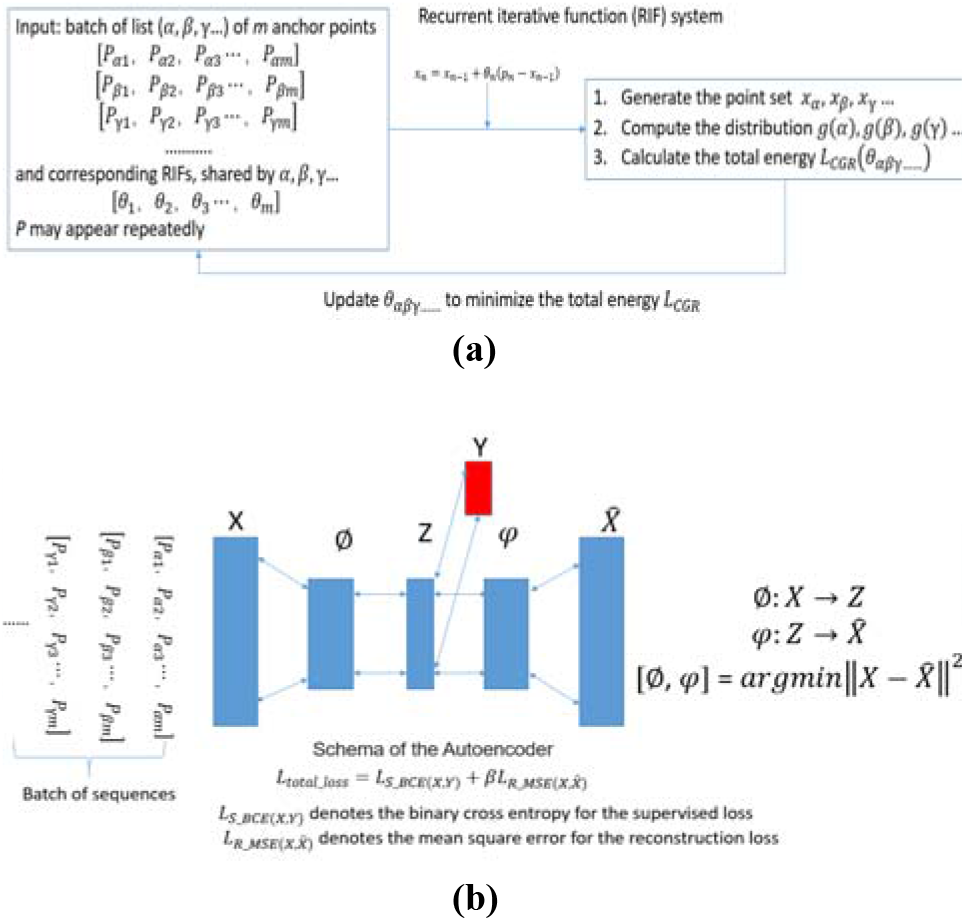
**(a) Algorithm A: *L_CGR_* minimization through optimization of SFs.** α, β, γ denote three amino acid sequences, *P_αi_* denotes the position of an anchor point i of sequences α, and the positions of all anchor points are denoted by the input vectors. After each iteration, the system updates *θ_k_* by 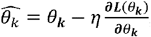, where *η* is the step rate. When RIFs converge (e.g., less than a predefined tolerance value), *L_CGR_* reaches the minimum and the iteration stops. **(b) Algorithm B:** SAE training. Ø and *φ* denote the encoder and decoder, respectively. X is a vector containing the positions of the anchor points from the corresponding sequences, and Y is the corresponding class with one-hot encoding. 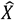 is the output of decoder *φ*, and the mean square error of the reconstruction loss for 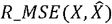 is defined by 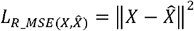.

### 2.2. Supervised Autoencoder (SAE)

SAE is known as an effective strategy with improved generalization capability to extract abstract representation of the input data (Liu et al. 2017; Le, Patterson, and White 2018), with addition of a supervised loss to the autoencoder in the representation layer, as shown in **Fig. 2(b)**. Let *X* denote the input vector which stores the positions of the anchor points obtained from the original sequences, *Y* denote the corresponding class, *L*_*S_BCE*(*X,Y*)_ be the supervised (primary) loss, and 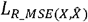 denote the loss for the reconstruction error:

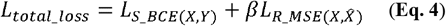

The supervised loss (*L*_*S_BCE*(*X,Y*)_) is to maximize the use of the known property labels to make the autoencoder more conducive to the final classification task. The second term represents the generalization capability of the decoder in reconstructing the data when using the encoded data from the latent space. After training, the encoder Ø (**Fig. 2b**) can be used to encode the original sequences into new sets of points in the latent space that improves classification. More importantly, these new point sets can be used to measure the distance between different sequences through calculating the Jensen-Shannon Divergence, and thus to further compute the corresponding *L_CGR_* of the whole system. While the first term (*L*_*S_BCE*(*X,Y*)_) is the primary loss, the coefficient *β* is a hyperparameter indirectly controls the relative strength of the supervised term (Siddhart et al. 2017). Based on our analysis, we found that the effect of optimization of *L_CGR_* is equivalent to minimizing *L_total_loss_*, as described next. If *β* = 0, SAE will degenerate into a multi-layer perceptron. In this situation, even if the solutions may well fit the data, the models will not find the underlying patterns in the data and the generalization will be poor. Therefore, we let *β* > 0 so as to improve the generalization performance of SAE. This is denoted as **Algorithm B** (**Fig. 2b)**.

To demonstrate the modeling process, we used 50 peptides from the databases described below and performed minimization of the total energy *L_CGR_* or *L_total_loss_*. The energy *L_CGR_* or *L_total_loss_* as a function of the training epochs are shown in **Fig. 3** (a, b, c, d, e, f). **Fig. 3(a, b)** illustrates the trajectories of *L_CGR_* minimization using **Algorithm A** with 3-mer motifs and MERCI motifs, respectively. **Fig. 3c** shows the trajectory of the total loss *L_total_loss_* with **Algorithm B**. **Fig. 3d** shows the effect of different β values on the convergence of *L_total_loss_*, and it demonstrates that the lower *β* leads to the faster SAE energy convergence. Similarly, **Fig. 3e** shows the trajectory of *L_CGR_* minimization using **Algorithm B**, with *β* = 0, based on MERCI motifs, while **Fig. 3f** represents the trajectories of *L_CGR_* minimization using **Algorithm B** with different *β*.

**Fig. 3.**
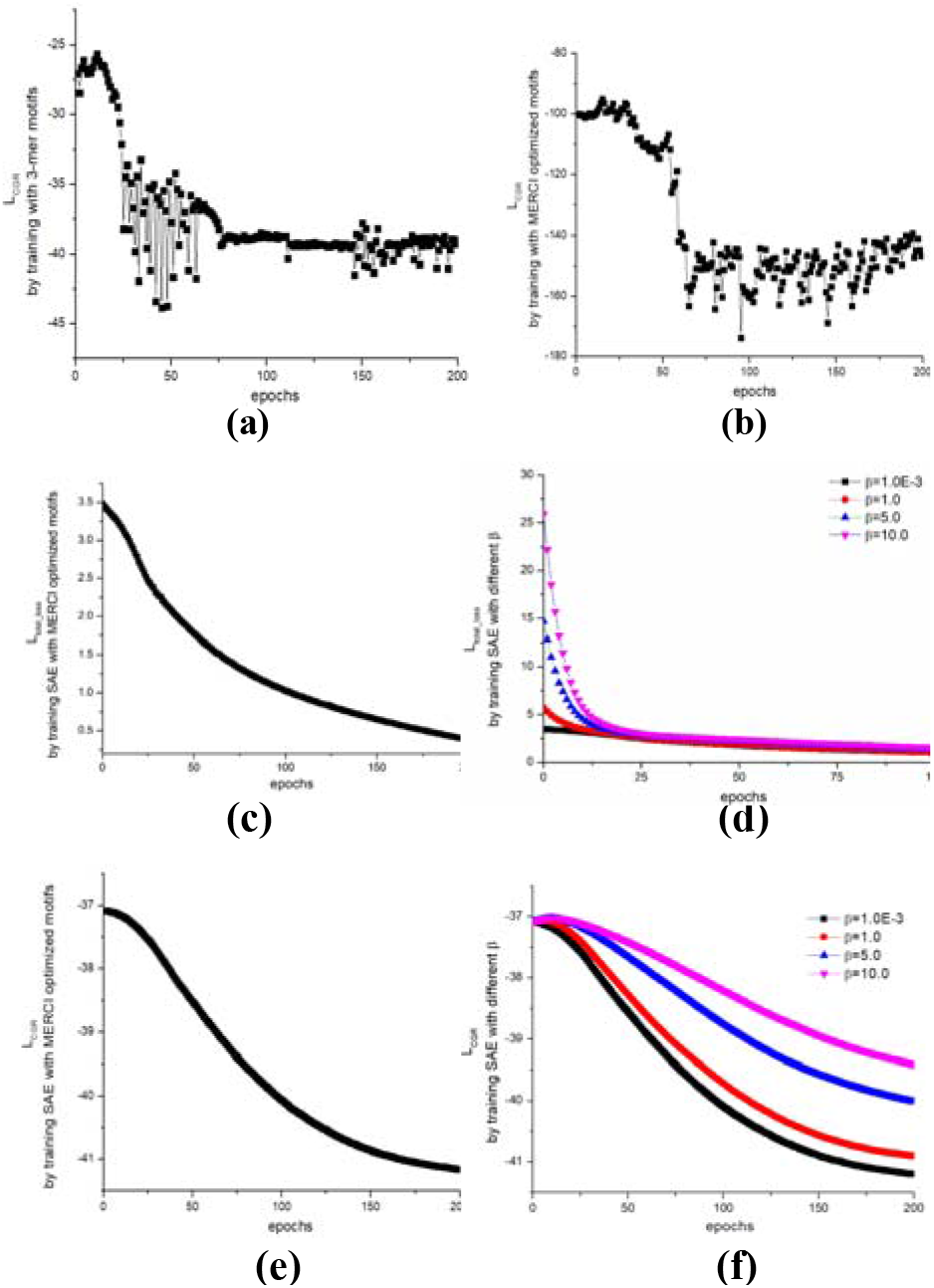
The trajectories of *L_CGR_* minimization using **Algorithm A** with 3-mer motifs (**a**) and MERCI motifs (**b**). We collected 50 peptides from HemoPI-1 Datasets, including 25 negative (non-hemolytic) and 25 positive (hemolytic) peptides. The step rates for **a,b** were set 1E-6 and 1E-4, respectively. (**c, d**) The trajectories of the total loss *L_total_loss_* with **Algorithm B** with the same input of motifs as described in (**a, b**), respectively, while d shows the effect of β on the energy convergence. (**e**). The trajectory of *L_CGR_* minimization using **Algorithm B**, with *β* = 0, using MERCI motifs. (**f**). The trajectories of *L_CGR_* minimization using **Algorithm B**, with different *β*, using MERCI motifs. It indicates that a lower *β* leads to faster convergence of the SAE model. (**g, h**) are two RGB images generated for the peptide “TKTTRNSPDSISIP” based on the SAE models at epoch=0 and 200, respectively, as illustrated in the trajectory in (**e**).

Interestingly, the above data shows that the minimization trajectories of both methods exhibit the same trend. Both and *L_CGR_* showed the similar monotonic decline, and this suggests that *L_total_loss_* can be used as an indicator for *L_CGR_* in terms of optimization to represent the original peptide sequences in the latent space. Therefore, we will use *L_total_loss_* instead of *L_CGR_* as a stop criterion for optimization process in **Algorithm B**. To further evaluate this concept, we applied the method to model our collections of hemolytic peptides and the HIV mutants. We also compared the performance of our method with 14 other binary classifiers, as described next.

## 3 Results and discussions

### 3.1. Datasets

#### Hemolytic Peptides

For hemolytic property modeling, we curated the data primarily from HemoPI-1, HemoPI-2, and HemoPI-3 (Chaudhary et al. 2016). HemoPI-1 contains 552 experimentally validated hemolytic peptides. However, the 552 non-hemolytic peptides are randomly generated, and thus they are just hypothetic negative data. HemoPI-2 includes 552 and 462 known hemolytic and non-hemolytic peptides, respectively. HemoPI-3 dataset is a bit larger, with 885 experimentally validated hemolytic peptides and 738 known low/non-hemolytic peptides. After combining all data and removing the duplicates, we obtained 1,150 experimentally confirmed hemolytic peptides and 1035 know non-hemolytic peptides, resulting in total of 2,185 sequences as shown in **Table. 1**. These peptides were used to build the final model, termed PepHemo, which has been deployed on our web server, as described in the next section, for public access to conduct in silico prediction of peptide hemolytic properties.

**Table 1.** Hemolytic Peptide Datasets.

|  | Training Sets |  | Validation Sets |  |
| --- | --- | --- | --- | --- |
|  | Pos | Neg | Pos | Neg |
| **HemoPI-1** | 442 | 442 | 110 | 110 |
| **HemoPI-2** | 442 | 370 | 110 | 92 |
| **HemoPI-3** | 708 | 590 | 177 | 148 |
| **Combined** | 868 | 813 | 272 | 222 |
| **Total** | 2185 |  |  |  |

#### HIVP Mutants

For HIVP sequences, the data were obtained for 5 FDA-approved protease inhibitors: indinavir (IDV), saquinavir (SQV), nelfinavir (NFV), amprenavir (APV), and lopinavir (LPV). The respective numbers of mutant protease sequences tested against each drug with their susceptibility are listed in **Table. 2**, as reported previously in (Rhee et al. 2006). Upon analysis of the combined data, we found that in total 776 unique HIVPs were tested against at least one drug, and 437 were tested against all 5 approved inhibitors. The data were used to construct models for predicting the susceptibility/resistance of HIVP mutants to each drug, namely **HIVP-SAE** which been deployed on our web server as described later.

**Table 2.** HIVP Datasets.

|  | IDV | SQV | NFV | APV | LPV |
| --- | --- | --- | --- | --- | --- |
| **Susceptible** | 384 | 457 | 303 | 424 | 223 |
| **Resistant** | 374 | 304 | 472 | 278 | 278 |
| **Total** | 758 | 761 | 775 | 702 | 501 |

Both datasets were then used to assess our **CGR-SAE** approaches. To this end, we adopted label-based metrics for binary classification and evaluated the derived models based on their confusion matrices using multiple parameters including accuracy, precision, and Receiver Operator Characteristic (ROC) curve as in **Appendix 3** in **Supplementary Materials**.

### 3.2. Classification of Hemolytic Peptides

We used our method on three datasets of the peptides to conduct classification based on their hemolytic activities. The overall workflow is illustrated in **Fig. S1**, and it primarily consists of two steps:

1. **Construct SAE models**. Each dataset was split into training and validation set as in **Table 1**. Starting with the peptide sequences in the training set, as illustrated in **Fig. 1** and **Fig. 2**, we first converted the sequences to the corresponding CGR data points. At this step, we also conducted cluster analysis. AS shown in **Fig S2**, all points are mixed together, indicating the classification task is challenging. Hence, we decided to build SAE encoder models through optimizing and SFs using **Algorithm B**. After such training, we obtained the optimized encoder models, which were used to generate (or encode) the 2D image representation for all the peptide sequences in both training and validation sets.
2. **Build convolutional neural network (CNN) classifiers**. Using the newly encoded 2D images for each peptide as described above, we performed another step of CNN to build classification models for the peptides. To this end, we performed 10-fold cross-validations to train the CNN models based on the training sets and evaluated the model performance based on the receiver operating characteristic (ROC) scores on the holdout validation datasets, and then select the best classifiers with the largest ROC area under the curve (AUC). We will also calculate the corresponding accuracy and precision for each classification models.

**Fig. 4a** illustrates the trajectories of the total cost function loss during model building with the training datasets. We selected the encoder for each dataset at epoch=200 where converges. With the derived encoder, each peptide sequence from both training and validation sets was transformed to a 2D image, and these encoded images were used to build convolutional neural network (CNN) binary classifiers based on the training sets and validated with the validation sets. To evaluate these SAE-based classification models, we calculated and plotted the model accuracy and precision as shown in **Fig. 4 (b, c)**. The ROC curve is shown in **Fig. 4(d)** which illustrates that our method is significantly better than the random way. We also compared our method with 14 other classifiers. For a fair comparison, we used the exactly same datasets as our input with the exact same splits of the training and validation sets as previously reported (Plisson, Ramirez-Sanchez, and Martinez-Hernandez 2020). The results are listed in **Table S1-3**, demonstrating that our CGR-based SAE approach can achieve the best classification with accuracy as high as 99%, or at least 80% for some challenging data. Overall, our models are superior to other methods (5-10% improvement), or at least comparable to a few such as GBC and XGBC.

**Fig. 4.**
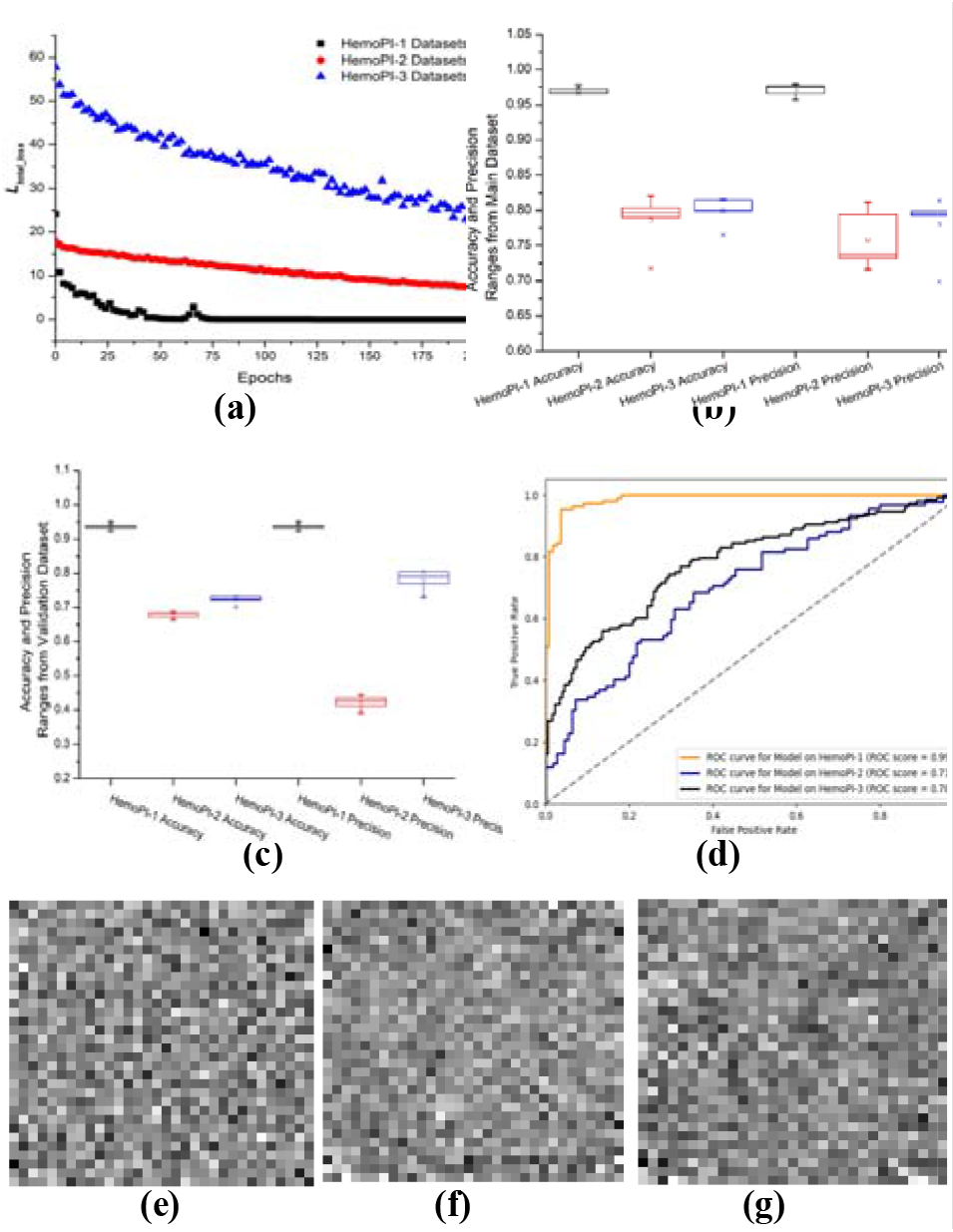
The trajectories of SAE training for three hemolytic peptide datasets, respectively. The sequences are represented with 30 CGR points based on a dictionary of 2,588 top ranked (highest frequency) MERCI motifs. For each dataset, the encoder at epoch=200 was selected. Then the encoded peptide images for each dataset were used to train our CNN classifiers. The classification accuracy and precision for respective training datasets are shown in **(b)**, and in **(c)** for the respective validation datasets. The colors are coded for different datasets. The ROC curves as shown in **(d)** were used to evaluate model performance. Using one of our models, we obtained the grayscale images with size of 32×32 for three peptides (GTPCGESCVYIPCISGVIGCSCTDKVCY LN; TCYCRRRFCVCVGR; SKHWLWLW), as illustrated in (**e, f, g**).

Of note, our method optimizes the classification task based on the distance distribution of the encoded 2D images, which can be characterized by the feature maps. We assume that the feature maps are similar if the images (i.e., peptides) belong to the same class, i.e., hemolytic or non-hemolytic. It has been reported that these feature maps in the discriminative image region can be used by the classifiers to perform the classification tasks, and they are termed as Class Activation Maps (CAM) (Zhou et al. 2016). To further interpret our results and demonstrate how our SAE approach could achieve accurate predictions, we selected 4 peptides, PC147 (hemolytic), PC148 (non-hemolytic), PC149 (non-hemolytic), and PC150 (hemolytic), all of which belong to a family of potent cationic antimicrobial peptides (Langham et al. 2008). **Fig. 5** shows the different representation of these peptides in CGR representations (3rd column) and SAE encoded 2D images (4th column). We computed the pair-wise distances which represent the similarity between sequences using the CGR representations as well as the encoded images, as defined in **Appendix 2.** As illustrated in **Fig. 5**, the only difference among the four peptides is at the 10th amino acid (the dashed rectangle in the 2nd column), and thus it is difficult to determine which peptide is hemolytic and which is non-hemolytic. Once converted to CRG representations (the 3rd column), the difference of these peptide sequences is amplified, but it is still a challenge for classification. Upon encoded into 2D images by SAE, the representation becomes highly complex (the 4th column) and difficult to understand with our naked eyes, although the patterns of the images are indeed different. This is not unexpected, as it is what a black-box alike deep learning method does. Then we derived the discriminative CAM (the 5th column) for each sequence and overlapped it with the SAE-encoded 2D image. This was used to highlight the important regions that were crucial to achieve the best classification. After this transformation, as shown in the 6th column, we obtained visually simplified, distinct heatmaps. As illustrated in **Fig. 5**, the hemolytic peptides (e.g., PC147 and PC150) have similar highlighted key blocks from the feature maps, and we expect that these features are crucial for the hemolytic activities of the peptides. Based on the results, it is apparent that the non-hemolytic peptides (e.g., PC148 and PC149) show quite different patterns, as demonstrated in both CAM (5^th^ column) and heatmaps (6^th^ column). It is also wroth mentioning that the minor difference of the peptides at the sequence level (only one amino acid) can be amplified by our CGR-based deep learning encoders, and thus significantly improving the classification accuracy, consistent to our initial hypothesis as described earlier.

**Fig. 5.**
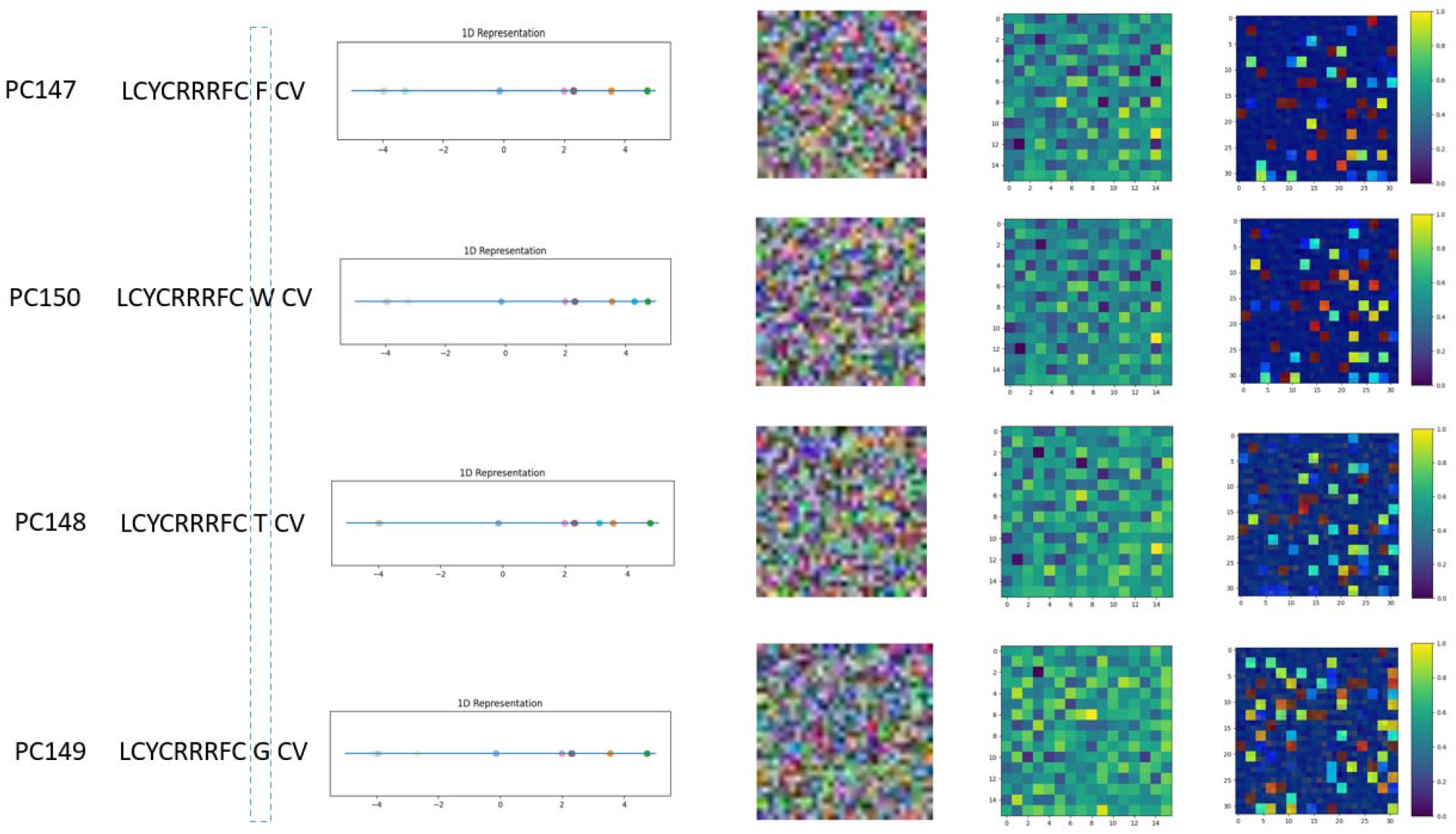
Comparative analysis of four peptides. Among them PC147 and PC150 are hemolytic; PC148 and PC149 are non-hemolytic. Their only difference is the 10^th^ amino acid, which is marked by the dashed rectangle in the 2^nd^ column. The 3^rd^ column is the CGR (1D) representation of the sequences. The 4^th^ column illustrates the SAE encoded 2D RGB images. The 6^th^ column are the heat maps, which were derived by overlapping the 2D encoded images (4th column) and the combined weighted convolutional feature maps, a.k.a., Class Activation Maps (CAM), as illustrated in the 5 ^th^ column obtained from the trained CNN model. Heat maps highlight the key blocks in feature maps (5 ^th^ column) and also show the distribution of the crucial pixels in the encoded images (4^th^ column).

### 3.3. Classification of HIVP mutants based on their susceptibility to approved drugs

We have long been interested in studying HIVP mutations, structures, and functions (Zhang, Kaplan, and Tropsha 2008). Because of its high mutation rate, the virus can rapidly evolve drug resistance (Rambaut et al. 2004). At the same time, the highly similar homologous sequences of HIVP mutants pose challenges to modeling studies such as binary classification. To further demonstrate the capability of our CGR-based deep learning approach, we employed the method to build models to predict whether a strain of HIVP mutant virus is resistant or susceptible to FDA approved HIVP inhibitors, as described earlier: APV, IDV, SQV, NFV, and LPV. Similar to classification of hemolytic peptides, we first constructed the SAE models to encode each HIVP mutant sequences, and then built CNN classifiers to predict whether a mutant is susceptible or resistant to a drug. Herein, we had to build such models for each drug.

**Fig. 6(a)** illustrates the trajectories of the total cost function loss (L_total_loss_) during the training of the HIVP SAE models for the 5 drugs based on the data in **Table. 2.** As shown, *L_total_loss_*, converges for all drugs at epoch=300, thus we used the SAEs at this point to encode HIPV sequences to derive the corresponding 2D images as their new representations, which were used for the second step of CNN training and model derivation. Such models can be used to select the most effective treatment strategies for patients based on their individual HIVP mutation profiles. The performance of the 5 selected models for respective drugs are illustrated with their ROC scores in **Fig. 6(b)**. The corresponding accuracy and precision of the respective training datasets and testing datasets are shown in **Fig. 6(c, d)**. Impressively, we have always achieved >95% (actually 100% for SQV) classification accuracies for both training and validation sets, demonstrating the significantly high predictive power of our approach. This indeed makes it possible and realistic to identify effective treatment options for patients based on their individual HIV protease mutation profiles.

**Fig. 6.**
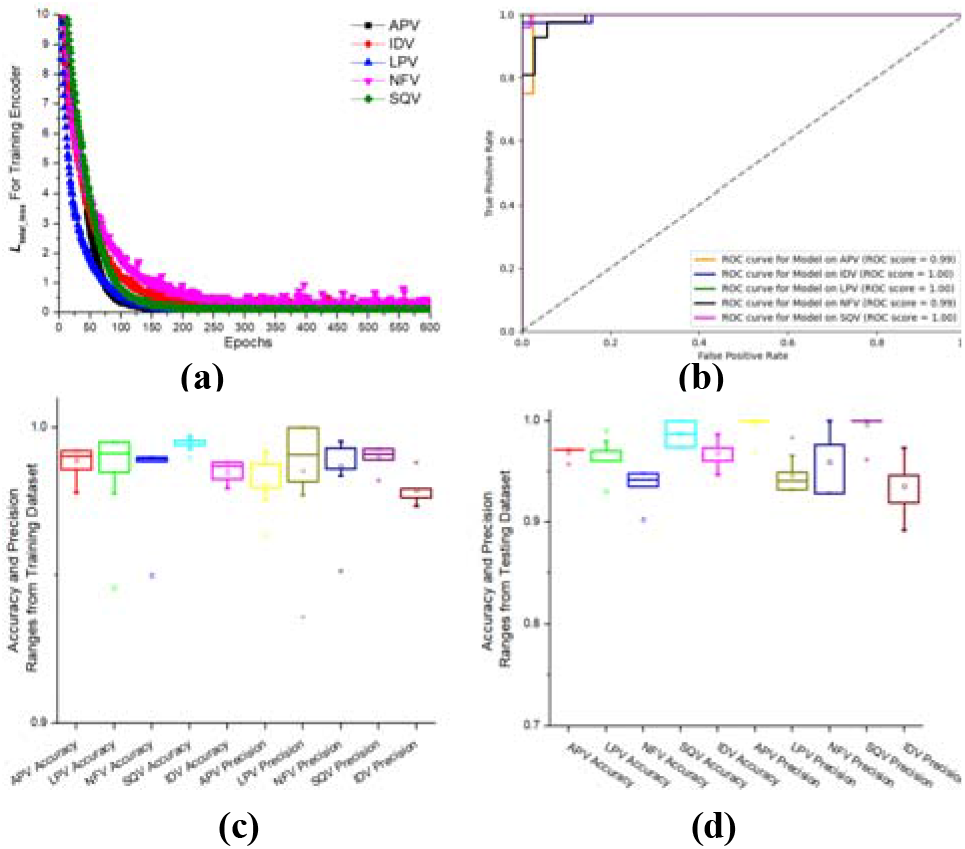
**(a)** The trajectories of cost function loss of SAE models trained with HIVP mutant datasets. The SAEs were trained with the HIVP mutant datasets tested on 5 drugs: IDV, LPV, NFV, SQV, APV. The converged SAE models were selected at epochs = 300 as the encoders to transform the protein sequences into 2D images. The CNN training was achieved using 10-fold cross-validations, and evaluated with the validation datasets. **(b)**. The ROC curves for the 5 datasets with their ROC scores, each corresponding to the best classification model for the respective drug. **(c)** and **(d)** show the classification accuracy and precision for respective training datasets and testing datasets, respectively.

However, for LPV and NFV, we were not able to accurately classify some HIVP mutants in terms of their susceptibility to the respective drugs (24 for NFV and 43 for LPV). There are a variety of reasons. As shown in **Fig S3**, the susceptible and resistant mutants are indistinctly mixed together based on our cluster analysis. Many of those mutants that were incorrectly classified are outliers to their classes. For instance, the mutant pos278 is susceptible to NFV (thus shown as a red dot), but it is far away from the other susceptible HIVPs (red dots), actually almost in the center of resistant proteases to NFV (blue dots, **Fig S3a**). This suggests that pos278 is more like an outlier to the susceptible HIVP mutants, and consequently our model was not able to correctly classify it into the right group. We had similar observations for this sequence in its susceptibility to LPV (**Fig S3b**), where experimentally it is susceptible to the drug (red dots), but it was clustered together with the resistant mutants and our modes could not accurately predict its susceptibility to LPV. Another reason of low accuracy for LPV predictions is probably that there are ~30% less experimental data available to LPV than other drugs (~500 vs. ~750 in **Table 2**).

To further understand from the biological point of view why we had false classifications for some HIVP mutant-drug pairs, we aligned 4 most frequently mis-classified mutants with the wild type HIVP (**Fig. S4**). Among them, three mutants (P118, P255 and P278) are all susceptible to NFV and LPV, but our models showed they are resistant to both drugs. The other mutant, N5, is resistant to IDV, but our model predicted the opposite. Based on their comparison with the wild type sequence, the most frequent mutations occur at L10, L63, I84 and L90. Other mutations can occur at K20, M36, M46, I54, A71, and V82. Indeed, it has been reported that L10 and L63 are frequent mutations leading to drug resistance (Flor-Parra et al. 2011; Appadurai and Senapati 2016; Natap 1998), and this was the pattern learned by our models. Surprisingly, the mutant P118 are still susceptible to NFV or LPV. Thus, our models could not predict this correctly. Another two mutants P255 and P278 even have more resistance-leading mutations including frequent mutations such as I84 and L90, along with K20, M36, or I54. Our model could not classify them correctly and it is not clear why they are still susceptible to the two drugs, but we think probably due to compensatory effects of multiple mutations (Piana, Carloni, and Rothlisberger 2002). For IDV, we have obtained a highly predictive model, with ~99% accuracy and ROC AUC is almost 1. Only two mutants were incorrectly classified, and N5 is one of them. By looking at the sequence, it does have frequent mutations such as I10, L63 and I84, thus it is not surprising that the mutant is resistant to IDV. The uniqueness of this sequence is that it has Leu insertion at position 10 and Val insertion at position 76. Probably such insertion mutations change the motif composition for this sequence, and thus our model could not characterize it correctly, as illustrated in **Fig S3c** where it was clustered within the susceptible mutants.

### 3.4. Web server implementation

Since it is labor-intensive, time-consuming, and costly to experimentally determine the hemolytic activities of a large number of peptides or test the drug susceptibility of HIV virus, we decided to share our highly predictive models via user-friendly web-based applications, namely PepHemo and HIVP-SAE, respectively. This provides the community to conduct in silico screening in an easy-to-access, ultrafast manner. As described earlier, the respective models were developed based on the CGR representation of amino acid sequences, followed by two steps of SAE encoding and CNN training. In addition, by combining all the reported datasets together, we curated the largest datasets to date for both projects to build the final respective classification models.

The web servers were implemented based on Python 3.8.10, torch 1.11.0, and Flask 2.4 web framework. Both PepHemo and HIVP-SAE have been designed and developed in a similar manner, with web page linked to each other. The interfaces have also been optimized for friendly mobile browsing, and primarily consist of four sections: Introduction, Input, Output, and Frequently Asked Questions (FAQ). The amino acid sequences (e.g., peptides, proteins) can be directly input to the online form in the FASTA format or uploaded as a file containing a list of FASTA sequences. In the Output section, the predicted results are provided in a table. A detailed instruction to data interpretation and the use of the programs are also provided on the web servers. Currently PepHemo is freely available at https://ai.imdlab.net/CGR_app/PepHemo and HIVP-SAE at https://ai.imdlab.net/CGR_app/HIVP_SAE.

## Conclusion

We adopted a SAE-based neural network to optimize the CGR representation of amino acid sequences (peptides and proteins). We first analyzed the relation between the optimization of CGR and the SAE-based encoder, and revealed that the optimized CGR in terms of SFs is equivalent to the corresponding SAE models. Based on this observation, we developed an optimized CGR-based SAE approach to encode amino acid sequences into 2D images, and such new representations are more conducive to more accurate classifications. To assess the effectiveness of our method, we first performed classification of hemolytic/non-hemolytic peptides based on the largest dataset we curated. We further employed the approach to classify longer amino acid sequences, i.e., mutant HIVP, for their susceptibility to FDA approved drugs. In both cases, compared with other classifiers, our models achieved superior performance, demonstrating that this method is efficient and universal. In the future, we will adopt the method to other classification tasks including multi-label, multi-type classifications which have been considered as more challenging problems. Additionally, we have built user-friendly web servers, PepHemo and HIVP-SAE, respectively, to share with the community so that others can further evaluate the method and use our models for their own biomedical discoveries.

## CONFLICTS OF INTEREST

There are no other conflicts to declare.

## FUNDING

BH was in part supported by NIH/NCI 1R01CA266187-01A1, 1R01CA225955-01, DOD W81XWH-20-PCRP-IDA, and Koch Foundation Grant.

## ACKNOWLEDGEMENTS

The authors are grateful to Drs. Cristina Ivan and Lon W. Fong for their helpful discussions and proofread of the manuscript.

## Notes

https://ai.imdlab.net:4443/HIVP_SAE

https://ai.imdlab.net:4443/PepHemo

### Competing Interest Statement

The authors have declared no competing interest.

### Summary of Updates

Update the affiliation of the coauthor

