## Supplementary Materials for "Sequence-based Optimized Chaos Game Representation and Deep Learning for Peptide/Protein Classification"

**Appendix 1.**

We set M points (number of top ranked motifs in the motif dictionary) randomly distribute on 1D x-axis as the anchor points denoted as $P_{k}$, $k=0,1,2\ldots M$. Based on these anchor points, we derive a sequence of encoded points $\left\{ X| x_{0},x_{1}, x_{2}\ldots\ldots x_{n} \right\}$, where *n* is the number of motifs in the sequence. Then we denote the origin at $x_{O}$ to define the spatial distribution of the encoded points by the distribution function$g(x)$,

$$g(x_{i})=\left| x_{i}-x_{O} \right|$$

where $\left| x_{i}-x_{O} \right|$ is the distance between point $x_{i}$ and $x_{O}$.

**Appendix 2.**

Assuming a sequence α is projected to a 2D matrix of size $m\times n$, and it can be represented as a 2D tensor, a greyscale or color image which is described in a 2D array $M\left[ x \right]\left[ y \right]$, where $x=0\ldots m,y=0\ldots n .$ The matrix can be flattened into 1D array and denoted by a point set $S_{\alpha}=\left\{ S_{\alpha}|x_{0}^{\alpha},x_{1}^{\alpha},x_{2}^{\alpha}\ldots x_{m\times n-1}^{\alpha} \right\}$. Similarly, we derive for a sequence β a point set $S_{\beta}=\left\{ S_{\beta}|x_{0}^{\beta},x_{1}^{\beta},x_{2}^{\beta}\ldots x_{m\times n-1}^{\beta} \right\}$.

Using the same distribution function introduced in **Appendix 1,** we denote $G\left( S_{\alpha} \right)$as the spatial distribution vector for the point sets $S_{\alpha}$ and $G\left( S_{\beta} \right)$ for the point sets $S_{\beta}$. Then the distance between the two sequences can be defined as

$D\left( \alpha,\beta\right)=JSD(G\left( S_{\alpha} \right),G\left( S_{\beta} \right))$

where $JSD$ function compute Jensen-Shannon Divergence:

$$JSD(G\left( S_{\alpha} \right),G\left( S_{\beta} \right))=\frac{KL\left( G\left( S_{\alpha} \right)\parallel G\left( M \right) \right)}{2}+\frac{KL\left( G\left( S_{\beta} \right)\parallel G\left( M \right) \right)}{2}$$

where $G\left( M \right)=\frac{1}{2}\left( G\left( S_{\alpha} \right)+G\left( S_{\beta} \right) \right)$, and $KL(G\left( S_{\alpha} \right)\parallel G\left( M \right))$ is Kullback–Leibler divergence that measure the distance between distribution of $G\left( S_{\alpha} \right)$ and $G\left( M \right)$.

**Appendix 3.**

We adopted label-based metrics (positive annotated as 1 and negative annotated as 0) for binary classification. The Accuracy, Precision, and Recall of dataset are defined as follows:

$\text{Accuracy}\text{ }\text{(}TP$, $FN$,$TN$, $FP\text{)}\text{ }\text{=}\frac{TP+TN}{TP+FP+TN+FN}$

$\text{Precision (}TP$, $FN$,$TN$, $FP\text{)=}\frac{TP}{TP+FP}$

where $TP$, $FN$,$TN$, $FP$ represent the type $j$ number of true positives, false negatives, true negatives and false positives, respectively [1]. The Macro-averaging general accuracy ACC, general precision PRE are defined as follows:

$$B_{Macro}\text{=}\frac{1}{q}\sum_{j=1}^{q} B(TP, FN, TN , FP)$$

where $B\in\left\{ \text{Accuracy, Precision} \right\}$, and the corresponding $B_{Macro}\in\left\{ \text{ACC, PRE} \right\}$, $q=2.$

**Table S1.** Comparison of PepHemo with 14 reported methods for classifying the hemolytic peptides in HemoPI-2. **Acc:** accuracy, **Prec:** precision

| **HemoPI-1** | **Model (442/442)** | | **Validation (110/110)** | |
| --- | --- | --- | --- | --- |
| **Classifiers** | **Acc** | **Prec** | **Acc** | **Prec** |
| LOGREG | 92.6 | 89.4 | 89.6 | 84.7 |
| KNN | 93.2 | 89.6 | 86.8 | 81.1 |
| CART | 90.4 | 85.8 | 83.7 | 77.6 |
| RFC | 93.8 | 91.1 | 87.3 | 81.8 |
| GBC | 94.0 | 91.3 | 90.4 | 86.5 |
| ADC | 93.8 | 90.7 | 91.4 | 87.4 |
| LDA | 94.2 | 91.1 | 90.5 | 87.1 |
| QDA | 91.5 | 88.3 | 85.5 | 83.8 |
| NB | 85.9 | 82.3 | 85.5 | 81.0 |
| SVC-LIN | 88.1 | 84.2 | 85.0 | 81.1 |
| SVC-RBF | 88.7 | 84.8 | 85.5 | 81.0 |
| SVC-POLY | 71.5 | 71.4 | 60.0 | 57.0 |
| SVC-SIG | 88.0 | 84.1 | 81.8 | 79.2 |
| XGBC | 94.8 | 93.3 | 92.4 | 86.7 |
| CGR-SAE | 99.4 | 98.9 | 94.1 | 97.3 |

**Table S2.** Comparison of PepHemo with 14 reported methods for classifying the hemolytic peptides in HemoPI-2 dataset. **Acc**: accuracy. **Prec**: precision

| **HemoPI-2** | **Model (442/442)** | | **Validation (110/110)** | |
| --- | --- | --- | --- | --- |
| **Classifiers** | **Acc** | **Prec** | **Acc** | **Prec** |
| LOGREG | 68.0 | 65.2 | 65.8 | 63.2 |
| KNN | 71.6 | 68.1 | 63.4 | 61.5 |
| CART | 69.6 | 67.0 | 62.4 | 61.9 |
| RFC | 69.0 | 66.0 | 63.9 | 62.1 |
| GBC | 76.7 | 72.9 | 72.3 | 68.9 |
| ADC | 71.3 | 68.1 | 73.8 | 70.2 |
| LDA | 70.0 | 66.6 | 59.9 | 59.7 |
| QDA | 66.4 | 64.5 | 60.9 | 60.5 |
| NB | 62.3 | 61.7 | 63.4 | 62.0 |
| SVC-LIN | 61.9 | 59.7 | 59.9 | 57.8 |
| SVC-RBF | 63.7 | 61.2 | 62.4 | 59.7 |
| SVC-POLY | 54.4 | 54.4 | 54.5 | 54.5 |
| SVC-SIG | 59.6 | 57.7 | 54.5 | 54.5 |
| XGBC | 76.1 | 70.3 | 69.3 | 67.2 |
| CGR-SAE | **80.9** | **75.4** | **67.9** | **67.8** |

**Table S3.** Comparison of PepHemo with 14 reported methods for classifying the hemolytic peptides in HemoPI-3 dataset. **Acc**: accuracy, **Prec:** precision

| **HemoPI-3** | **Model (442/442)** | | **Validation (110/110)** | |
| --- | --- | --- | --- | --- |
| **Classifiers** | **Acc** | **Prec** | **Acc** | **Prec** |
| LOGREG | 69.7 | 66.1 | 70.8 | 67 |
| KNN | 72.8 | 69.3 | 65.8 | 63.4 |
| CART | 68.9 | 66.6 | 64 | 62.7 |
| RFC | 72.4 | 68.3 | 66.8 | 64.1 |
| GBC | 75.8 | 71.9 | 72.9 | 69.4 |
| ADC | 72.3 | 68.6 | 71.4 | 68.1 |
| LDA | 70.8 | 66.9 | 70.8 | 67.3 |
| QDA | 65.9 | 64.0 | 59.4 | 59.1 |
| NB | 64.4 | 62.8 | 65.9 | 64.0 |
| SVC-LIN | 65.4 | 62.0 | 66.5 | 62.8 |
| SVC-RBF | 66.3 | 62.7 | 66.1 | 62.8 |
| SVC-POLY | 54.6 | 54.6 | 54.5 | 54.6 |
| SVC-SIG | 63.4 | 60.4 | 63.4 | 60.3 |
| XGBC | 74.7 | 74.3 | 73.2 | 67.6 |
| SAE-CNN | **81.9** | **76.5** | **73.8** | **77.7** |

**Fig. S1.** The overall workflow of CGR-based SAE and CNN model building. The initial amino acid sequences are converted to VGR data point based on a dictionary of motifs. They are then used as input to build SAE models to derive the encoders, which will be used to encode the sequences into 2D images. These images are hypothesized to better represent the sequences, and they are employed to build CNN models as classifiers which are further validated with the holdout test set. These final models were deployed on our web server.

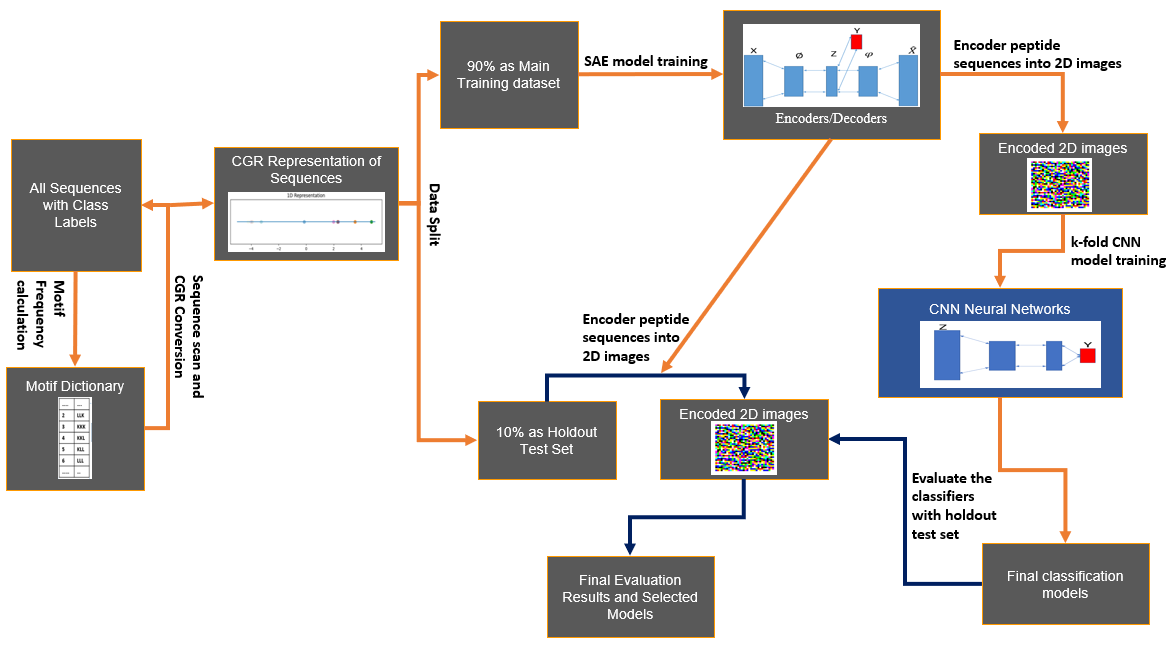

**Fig. S2.** Cluster analysis of all 2,185 hemolytic (red) and non-hemolytic (blue) peptides based on CGR data point representation generated using the ranked MERCI motif dictionary.

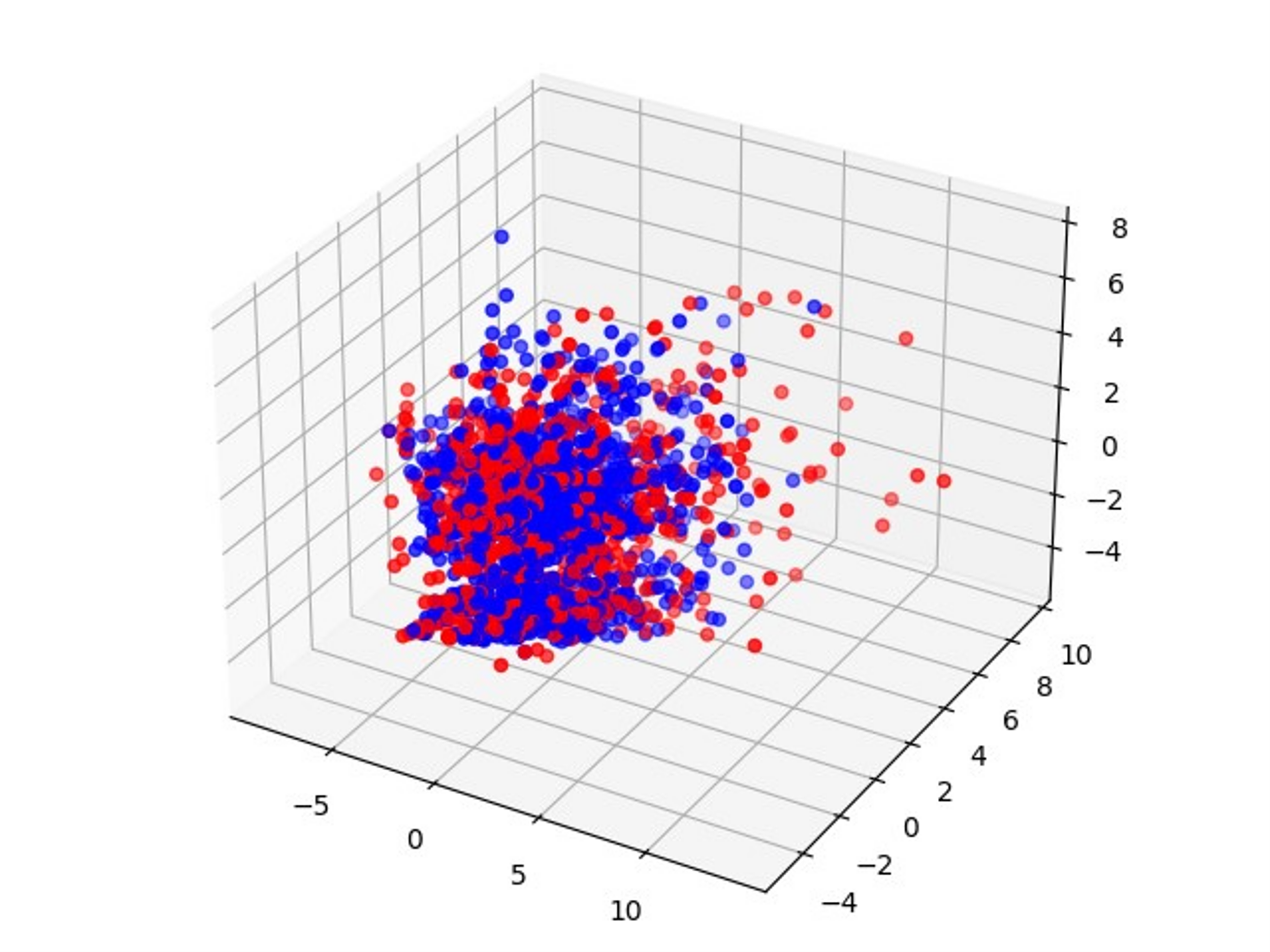

**(c).** Cluster Analysis of HIVP Mutants tested against IDV.

**(a).** Cluster Analysis of HIVP Mutants tested against NFV.

**(b).** Cluster Analysis of HIVP Mutants tested against LPV.

**Fig. S3.** Cluster analysis of all HIVP mutants that have been tested against **(a)** NFV, **(b)** LPV, and **(c)** IDV, respectively. The 3 principal components (PC1, PC2, and PC3) were derived based on the CGR representation of HIVP sequences. Red represents those HIVP mutants susceptible to the drugs, and blue for resistant sequences. The black-circled, enlarged dots represent: **(a)** Mutant P278: PQITLWQRPIVTIK VEGQLREALLDTGADDTVLEDINLPGKWKPKMIGGIGGFVKVKQYDNVHMEICGHRAIGTVLVGPTPVNIIGRNMMTKIGCTLNF. This mutant is an outlier: it is susceptible to NFV (thus in red), but it is far away from the major susceptible mutants (red). Instead, it is within the blue cluster (resistant to NFV), and it was incorrectly predicted as resistant to NFV. **(b)** The same mutant P278 is also susceptible to LPV (thus in red), but it is also far away from the major susceptible mutants (red). Instead, it is within the blue cluster (resistant to LPV), and it was incorrectly predicted as resistant to LPV. So, it is also an outlier to the LPV susceptible mutants. **(c)** Mutant N5: PQITLWQRPLVTIKIGGQLKEALLDTGADNT VLEEMNLPGRWKPKMIGGIGGFIKVRQYDQIPIEICGHKAIGTVLIGPTPVNIIGRDLMTQIGCTLNF. This mutant is also an outlier: it is resistant to IDV (thus in red), but it is far away from the major resistant mutant cluster (blue). Instead, it is within the red cluster (susceptible to IDV), and it was incorrectly predicted as susceptible to IDV.

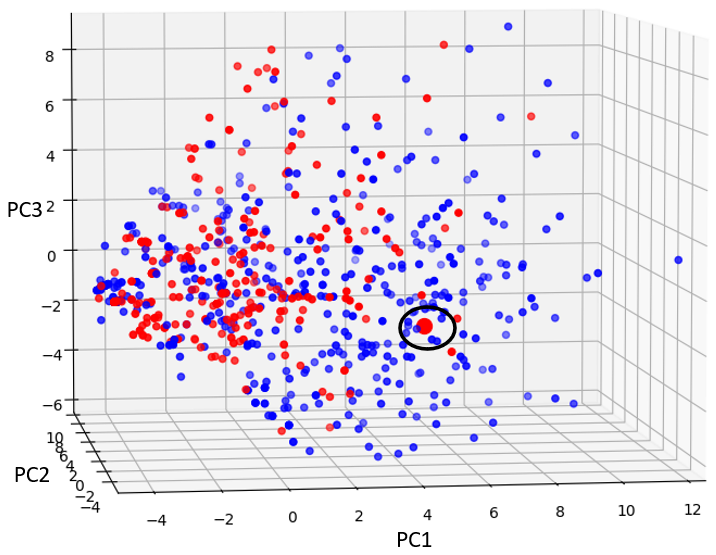

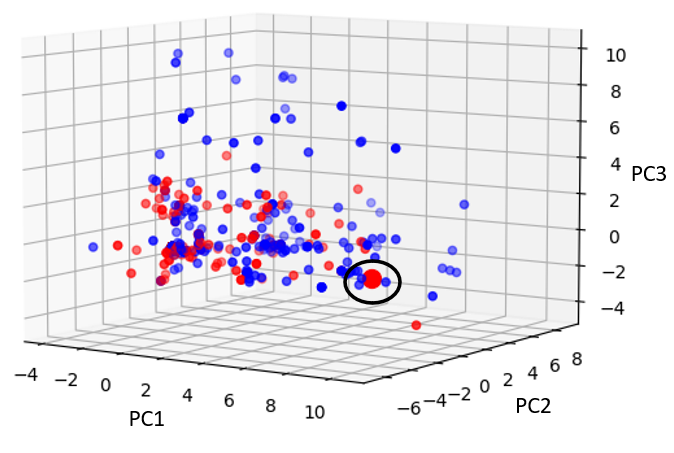

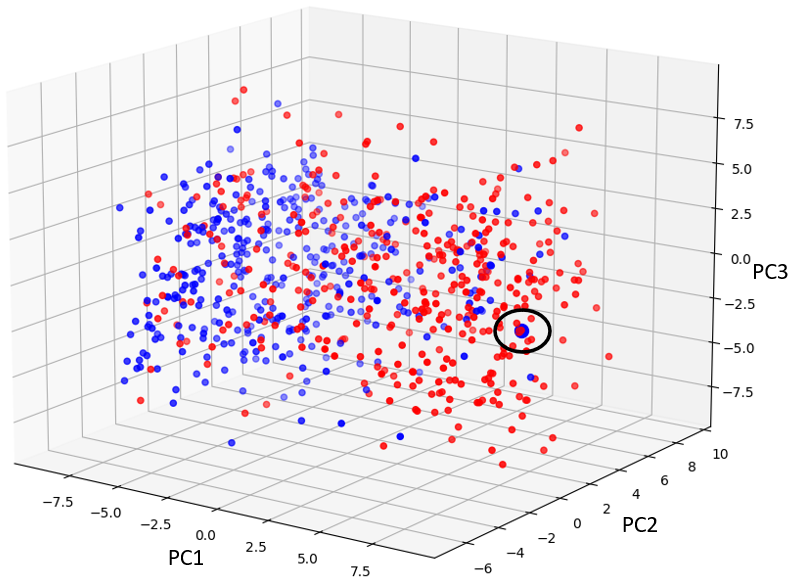

**Fig. S4.** Sequence alignment of 4 poorly predicted mutant sequences with the wild type HIVP. The green represents the identical residues in all 5 proteases, and the red represents the positions with mutations in at least one sequence. P118, P225 and P278 are susceptible to both NFV and LPV but incorrectly classified by our models for both drugs. N5 is resistant to IDV but our models predicted the opposite.

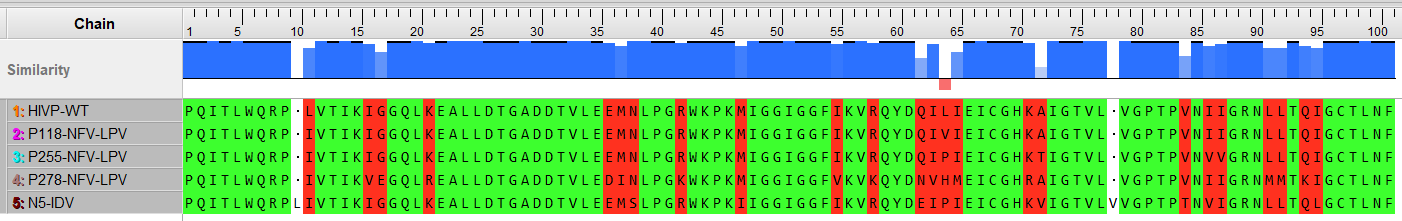
